# Cellular, Molecular, and Enzymatic Signatures of Thrombi are Vascular Bed-Dependent

**DOI:** 10.1101/2022.08.11.503688

**Authors:** Matthew Bender, Anu Aggarwal, Doran Mix, Matthew Godwin, Suman Guntupalli, Aravinda Nanjundappa, Leben Tefera, Ihab Hadadin, Rohan Bhandari, Michael Tong, William M. Baldwin, Robert L Fairchild, Marcelo Gomes, Joseph Campbell, David Schumick, Scott J. Cameron

## Abstract

**Background:** The contribution of arterial and venous thrombi to vascular remodeling is unclear. While catheter-extraction of thrombus in cerebrovascular accident (CVA) is time-sensitive, similar urgency is rare in managing venous thromboembolism (VTE).

**Objectives:** Our goal was to determine molecular cellular signatures of thrombus extracted by catheter from various vascular beds to gain insight into vascular remodeling.

**Methods:** Twenty-five patients underwent catheter-directed thrombectomy (CDT), 13 for acute CVA, 8 for pulmonary embolism (PE), and 4 for deep vein thrombosis (DVT). Protein and RNA extracted from thrombus was evaluated by immunoblotting and sequencing, respectively. Thrombus-derived enzymes for which substrate is present in the blood vessel wall were examined for enzymatic activity.

**Results:** Time from symptom onset to thrombus extraction was 7.7 ± 1.9 hours for CVA and 109 ± 55 hours for VTE. Protein concentration, white blood cell and red blood cell content were all greater in venous compared with arterial thrombus while platelet content was similar. Both venous and arterial thrombus contained multiple Matrix Metalloproteinase (MMP) isoforms. MMP9 specific activity was greater in venous than in arterial thrombus (57 ± 6 ng/mL.μg protein^-1^ vs. 24 ± 8 ng/mL.μg protein^-1^, P=0.0051).

**Conclusions:** Arterial and venous thrombus have dissimilar phenotypes, each with biologically-active enzymes known to remodel blood vessels, and enzymatic activity proportional to the white blood cell content which increases with thrombus age. These data suggest a mechanistically-important role for early CDT to avoid the consequences of irreversible vascular remodeling.

**Condensed Abstract:** Emergent extraction of acute thrombus from arterial vascular beds restores limb and end-organ perfusion and is widely-accepted to be the standard of care. Extraction of thrombus from venous vascular beds, however, is rarely considered urgent, even though many patients subsequently develop debilitating symptoms. By capitalizing on privileged thrombus extracted from multiple vascular beds, we gained mechanistic insight regarding the cellular composition and cell-derived enzymes secreted from thrombus that may remodel the vessel wall. This study shows thrombi are biologically-active entities, continuously recruiting circulating cells that secrete enzymes both proportional to thrombus age and the time of patient presentation.

## Introduction

Systemic anticoagulation for venous thromboembolism consisting of DVT or PE is the standard of care unless contraindicated (1). Acute PE is a true thrombotic emergency with cardiovascular mortality surpassed only by acute CVA and myocardial infarction (MI) (2,3). For patients with acute PE and high-risk features including right ventricle (RV) strain or cardiogenic shock, escalation of treatment beyond anticoagulation alone may include systemic thrombolysis or percutaneous procedures including catheter-directed thrombectomy (CDT) (4,5). Similarly, for patients with acute DVT, anticoagulation alone is the standard of care unless contraindicated. However, there is no clear role for additional therapies (6). Thrombectomy has been employed in the treatment armamentarium of select cases of acute DVT with mixed results. Phlegmasia cerelea dolens (PCD), a condition causing compromised of arterial inflow vessels secondary to a massive venous thrombus burden, remains the only true indication to employ CDT acutely (7).

While the consequences of acute arterial and venous thrombosis gain most attention in clinical studies, the long-term consequences of chronic thrombosis remain under-studied. Most acute thrombi in the pulmonary artery and the venous vasculature of the lower extremities gradually resolve over time when the patient is treated with therapeutic anticoagulation to prevent further thrombotic insults. However, registry data indicates incomplete thrombus resolution is a feature of more than 50% of patients following acute DVT or PE. The pathological processes behind incomplete thrombus resolution are incompletely understood (8). In the lower extremities, residual thrombus in deep veins may alter the integrity of the vein wall by causing incomplete valvular coaptation, venous hypertension, skin discoloration and ulceration which are features of the debilitating condition post-thrombotic syndrome (PTS) (9). Similarly, incomplete thrombus resolution in the pulmonary vasculature and vascular remodeling may lead to pulmonary hypertension, and occasionally the debilitating condition of chronic thromboembolic pulmonary disease (CTED) (10).

In patients with acute CVA, there is robust evidence demonstrating improved functional outcomes after endovascular thrombectomy (11–13). Prospective trials using CDT conflict on whether embolectomy improves functional outcomes in patients who sustain an acute DVT or PE raising some concern for routine off-label use of percutaneous devices for the purposes of thrombus retrieval (14–17). While there is little doubt regarding the debilitating impact of incomplete thrombus resolution on the functional status of patients following DVT, high-quality prospective studies have shown mixed efficacy for CDT (18–20).

Since thrombus retrieval is clearly beneficial in patients with acute CVA, an important consideration regarding the efficacy of CDT for thrombus extraction in the context of DVT and PE may be the timing of the intervention. Unlike an acute cerebrovascular insult which alerts patients immediately to functional deficits, patients with VTE sometimes have subtle symptoms for many days or weeks, allowing the aging thrombus to potentially initiate vascular remodeling. Patients with PTS commonly have lifestyle-altering symptoms that are occasionally debilitating. In a *posthoc* analysis of the ATTRACT trial, Li *et al*. demonstrated an apparent benefit to patients who underwent early CDT (within the first 48 hours), demonstrating mechanistically in a murine model of DVT that early restoration of blood flow prevented venous remodeling and fibrosis (21).

We previously observed that thrombus extracted from arteries of patients with acute myocardial infarction or from the wall of patients with chronic infrarenal abdominal aortic aneurysm (AAA) secretes activated enzymes, including those belonging to the zinc-dependent matrix metalloproteinase (MMP) family (22,23). Based on this data, we hypothesized that thrombus from any vascular bed may not be inert—as was previously believed—but rather may be biologically active, with unique molecular and cellular signatures based on its location. We therefore compared thrombus extracted from the lung in patients with acute PE to patients with thrombus extracted following acute DVT, comparing and contrasting this to properties of thrombus extracted from patients by catheter-directed arterial embolectomy following CVA.

## Methods

### Study approval

This study complies with the Declaration of Helsinki and was approved by the University of Rochester RSRB for the analysis of thrombus that would otherwise be biological waste from patients presenting to the neurological surgery service and the vascular surgery service. Each study participant provided written informed consent for utilization of tissue for biomedical research.

### Thrombus Gene Analysis

Arterial thrombi (n=6 infrarenal aorta, n=6 brain) and venous thrombi (n=5 leg, n=6 pulmonary artery) were evaluated for expression of genes related to fibrosis and inflammation. Thrombus extracted at the time of clinical intervention was snap frozen in liquid nitrogen and stored until future analysis. RNA was extracted from thawed thrombus using TRIzol (Thermofisher) after homogenizing the sample using TissueLyser II (QIAGEN) and then purifying the RNA (RNeasy mini kit, QIAGEN). For RNA sequencing, 100 μg RNA per sample was hybridized to a Panel Plus codeset consisting of nCounter ^®^ fibrosis v2 panel supplemented with 12 additional *probes:MMP20, MMP21, CD235a, CD69, GP1BA, GP1BB, ITGA2B, GP9, GP5, vWF, CXCL4*, and *CD62*. Data normalization and gene expression analysis used ROSALIND ® WITH A HYPERSCALE ARCHITECTURE using nSolver version 4.0.7 and with advanced analysis version 2.0.134 (ROSALIND inc., San Diego, CA). Adjusted p-values were used to determine the expression of each gene was quantified and pathways analysis was used to determine the biological systems involved in each thrombus.

### Thrombus MMP Analysis

#### Quantitative Polymerase chain reaction

a custom-designed MMP array (Bio-Rad) was employed using immobilized and lyophilized primers on a 96 well-plate for the following MMP isoforms: *MMP1, MMP2, MMP7, MMP9, MMP10, MMP14, MMP16, MMP20, MMP21* along with three housekeeping genes for increased confidence (*GAPDH, ACTB, ITGA2B*). For each sample, 100 ng RNA was used to prepare cDNA using High-Capacity RNA-to-cDNA™ kits (Applied Biosystems). qPCR was performed for each candidate gene in quadruplicate on CFX96 Touch Real-Time PCR Detection System (Bio-Rad) with itaq Universal SYBR Green Fast qPCR Mix. Data were analyzed by taking the mean ratio of Cq (Quantification Cycle) values of genes of interest to the housekeeping gene.

#### Gel Zymography

homogenized thrombus (7 mg) lysates were centrifuged for 15 mins at 4°C, and supernatants were placed in 50% volume/volume 2x non-reducing sample buffer at the following final concentration: Tris-HCl 250 mM, 0.5% SDS, 1% glycerol, 0.05% bromophenol blue for 10 minutes without boiling., then loaded onto a pre-cast gel with a gelatin matrix (10% zymogram, #ZY00102BOX, Invitrogen). The gel was renatured by gently rocking in 2.5% Triton-X-100 for 30 mins at room temperature, then allowed to equilibrate at room temperature with gentle rocking in zymogram buffer with the final concentrations: Tris-base 50mM, NaCl 0.2M, CaCl2 5mM, Tween-20 0.02% for 30 mins before decanting, and incubating in fresh zymogram buffer for 12 hours overnight at 37°C. The zymogram buffer was decanted, and the gel was rocked at room temperature for 4 hours in Simply Blue Safestain (Invitrogen). MMP activity was noted by clear bands in the final gel (a reverse image). Total MMP activity in each lane was quantified by densitometry using ImageJ software (NIH).

#### MMP Activity Assay

protein extracted from arterial and venous thrombus was quantified and 600 ug/mL was used per sample to determine MMP9 activity using a substrate specific chromogenic assay according to the manufacturer’s recommendation (cat. # F9M00, R & D Systems).

### Immunoblotting

Homogenized thrombus lysates were centrifuged for 15 minutes at 4°C, and supernatants were placed in 50% vol/vol 2x denaturing sample buffer at the following final concentration: Tris-HCl 250 mM, 0.5% SDS, 1% glycerol, 0.05% bromophenol blue, 5% β-mercaptoethanol for 10 minute. Protein lysate (40 mg per lane) was separated by SDS-PAGE (5-20% gradient, Bio-rad) at 125V (constant voltage, room temperature) on commercially-available gradient gels (4-20%, Invitrogen). Separated proteins were then transferred to nitrocellulose membranes (Bio-Rad) at105 V for 1 hour with an ice pack at room temperature. Blocking buffer was 3% bovine serum albumin/Tris-buffered saline–Tween 20 for 60 minutes at room temperature with agitation. TBST-T Primary antibody incubation (1:1000 dilution in 3% bovine serum albumin/Tris-buffered saline–Tween 20) for 12 hours at 4 degrees C with agitation. Secondary antibody (GE Healthcare, Buckinghamshire, UK) was used in a 1:2000 titer in 5% milk/Tris-buffered saline–Tween 20 for 1 hour at room temperature with agitation. Final autoradiographic films (Bioblot BXR, Laboratory Product Sales, Rochester NY) were quantified by densitometry using ImageJ software (National Institutes of Health)

### Bioinformatics and Statistical Analysis

RCC files from NanoString nCounter system were quality controlled and analyzed using nanoR package in R (https://github.com/KevinMenden/nanoR). Normalized count matrices were used for differential gene abundance using DeSeq2 package^1^. Differentially expressed genes were used to identify the functional enrichment using GO enrichment the clusterProfiler package^2^ (version 3.04) (24,25). Graphical data are shown as mean ± SEM unless stated. Equal variance between analysis groups were evaluated by the Shapiro-Wilk test. For normally distributed data between two groups, a 2-tailed Student’s *t* test was used. For nonparametric data, the Mann-Whitney *U* test was used. Significance was accepted as a *P* value of less than 0.05. Analyses were conducted using GraphPad Prism 7 (GraphPad Software).

## Results

### Patient demographics and procedural details

25 patients (13 acute CVA, 8 PE, 4 DVT) were enrolled in this study. The three groups did not differ by race or sex. Patients with stroke were slightly older than those with VTE (74 years vs. 62, p=0.0528), and with a lower BMI (24.0 Kg/m^2^ for CVA vs. 35.0 Kg/m^2^ for VTE, p=0.0006). Patients did not differ in relevant comorbidities or premorbid antiplatelet and anticoagulant use. The majority of patients with acute CVA received either intravenous or intra-arterial tPA (71% with CVA vs. 0% with VTE p=0.001). All patients with PE and DVT received systemic heparin while fewer patients with acute CVA who received full-dose IV tPA typically did not (50%, p=0.016). On follow up, 25% of patients with DVT had clinical symptoms consistent with post-thrombotic syndrome (PTS), while 16.7% of patients with acute PE had clinical symptoms consistent with CTED. (**Table 1**). Time from symptom onset to thrombus extraction was les for stroke patients (7.7 hours for stroke vs. 109 hours for VTE, p<0.0001) (**Figure 1A**).

**Fig. 1A.**
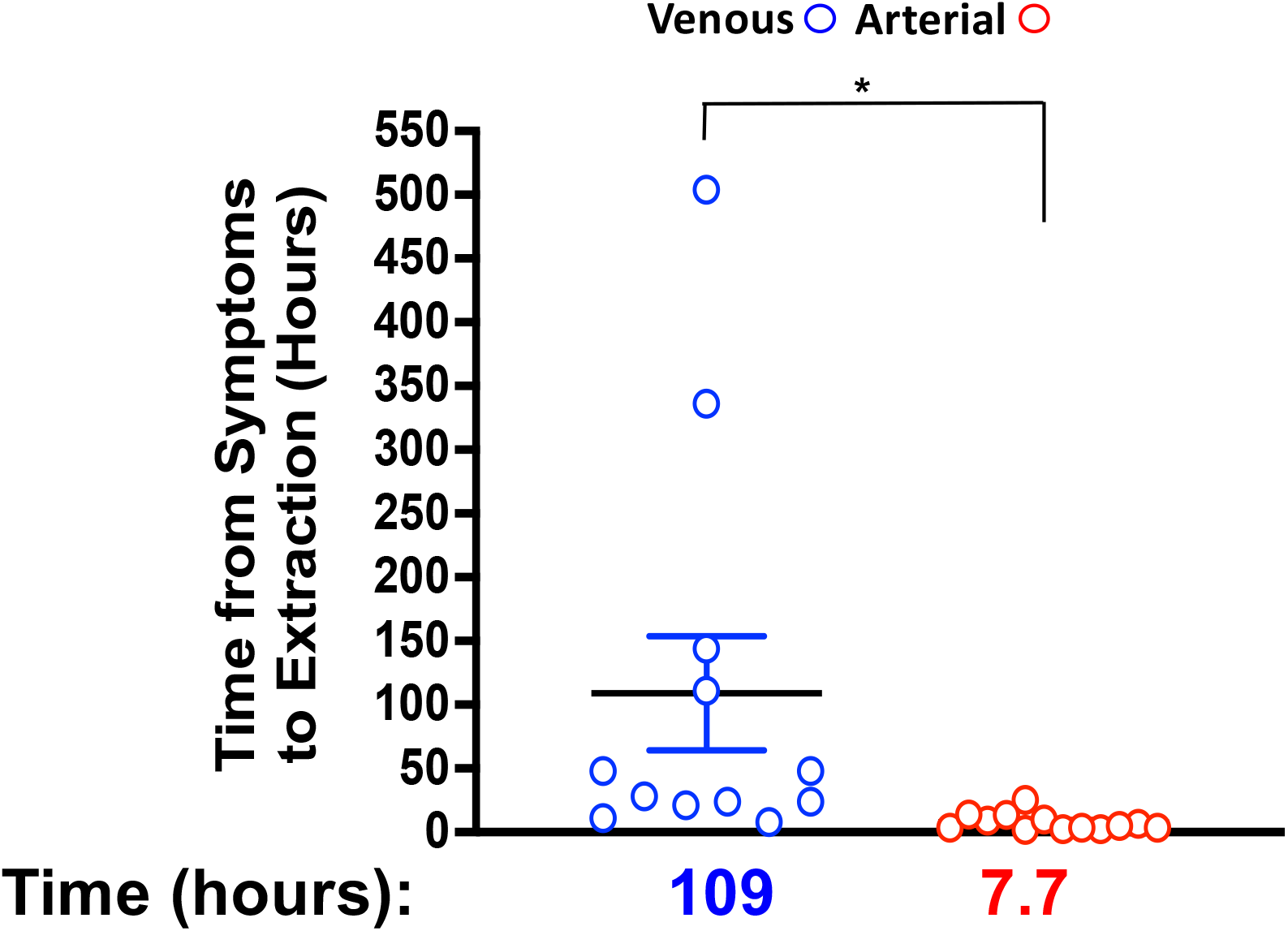
Time from symptoms onset to thrombus extraction. Time from symptom onset for acute thrombotic stroke (n=13) and VTE (DVT and PE, n=12) was reported as mean ± SEM. *P <0.0001 by Mann-Whitney *U*. VTE=venous thromboembolism.

**Table 1.** Characteristics of study population: Patients with acute thrombotic stroke and VTE (DVT and PE) were included in the study. Data are reported as mean ± SD unless otherwise stated with difference between groups as noted.

|  | Venous<br>thromboembolism<sup>†</sup><br>(N = 12) | Arterial<br>thromboembolism<sup>†</sup><br>(N = 13) | Triage (Extraction)<br>Time, mean hours (SD) | p-value<sup>†</sup> |
| --- | --- | --- | --- | --- |
| Age, mean (SD) | 62 ± 17 | 74 ± 11 | -- | 0.0528 |
| Male (%) | 6 (50.0%) | 5 (38.5%) | 64.3 (101.87) | -- |
| Female (%) | 6 (50.0%) | 8 (61.5%) | 50.0 (131.32) | -- |
| BMI, mean (SD) | 35 ± 9 | 24 ± 4 | -- | 0.0006 |
| **Race, n (%)** |  |  |  |  |
| Asian | 1 (8.3%) | 0 (0%) | 21 (0) | -- |
| Black | 2 (16.7%) | 2 (15.4%) | 15.2 (11.11) |  |
| White | 9 (75%) | 10 (76.9%) | 69.3 (132.37) |  |
| Unknown | 0 (0%) | 1 (7.7%) | 9.1 (0) |  |
| **Thrombus Type, n (%)** |  |  |  |  |
| Cerebrovascular | 0 (0%) | 13 (100%) | 7.7 (6.75) | -- |
| Deep Vein Thrombosis (DVT) | 4 (33.3%) | 0 (0%) | 47.75 (44.97) |  |
| Pulmonary Embolism (PE) | 8 (66.7%) | 0 (0%) | 139.5 (183.72) |  |
| Massive PE (SBP <90, syncope, Shock) | 5 (41.7%) | 0 (0%) | 88.8 (138.92) |  |
| Sub massive PE (RV strain) | 9 (75%) | 0 (0%) | 136 (172.36) |  |
| **Vessel Occluded, n (%)** |  |  |  |  |
| MCA | 0 (0%) | 8 (61.5%) | 8.4 (8.02) | -- |
| ICA | 0 (0%) | 4 (30.8%) | 6.08 (5.18) |  |
| Basilar | 0 (0%) | 1 (7.7%) | 9.1 (0) |  |
| RCIV | 1 (8.3%) | 0 (0%) | 111 (0) |  |
| LSFV | 1 (8.3%) | 0 (0%) | 48 (0) |  |
| LCIV | 2 (16.7%) | 0 (0%) | 16 (7.07) |  |
| Main PA, L/R Pas | 4 (33.3%) | 0 (0%) | 224 (237.05) |  |
| RV, Main PA | 1 (8.3%) | 0 (0%) | 24 (0) |  |
| Distal PA's | 1 (8.3%) | 0 (0%) | 28 (0) |  |
| L/R Pas | 1 (8.3%) | 0 (0%) | 24 (0) |  |
| 'R PA | 1 (8.3%) | 0 (0%) | 144 (0) |  |
| **Post-thrombotic syndrome?** |  |  |  |  |
| Yes | 3 (25%) | 0 (0%) | 22.3 (22.28) | -- |
| No | 6 (50%) | 0 (0%) | 174.67 (199.86) |  |
| Unknown | 3 (25%) | 0 (0%) | 64 (69.28) |  |
| **Post-PE Syndrome/CTED** |  |  |  |  |
| Yes | 2 (16.7%) | 0 (0%) | 266 (336.58) | 0.041 |
| No | 7 (58.3%) | 0 (0%) | 83.29 (116.83) |  |
| Unknown | 3 (25%) | 0 (0%) | 64 (69.28) |  |

<sup>†</sup>: denotes the data sets between which the p-value is calculated via t-test

### Thrombus Cellular Characteristics and Biological Activity

Older thrombi extracted from the venous circulation had had a higher protein concentration per 7 mg of dry tissue than thrombi extracted from the arterial circulation (3.1 mg/mL vs. 2.1 mg/mL, p=0.031) (**Figure 1B**). We previously demonstrated that arterial thrombus extracted from patients with MI or from patients with infrarenal abdominal aortic aneurysms (AAA) are enriched in activated MMPs (22,23). As such, a bespoke MMP gene array was utilized to evaluate the presence of MMPs in arterial and venous thrombi. The quantity of MMP isoforms was similar in arterial and venous thrombus with the notable exception that MMP7 and MMP21 were not detected in arterial thrombus. Genes for the following MMP isoforms were amplified in both arterial and venous thrombus in every patient: MMP1, MMP2, MMP8, MMP9, MMP10, and MMP14 (**Figure 2A**). Following the separation of protein extracted from venous and arterial thrombus by SDS-PAGE, in gel zymography revealed a very strong signal for an activated MMP isoform around 95 KDa corresponding to the mass of MMP9 (**Figure 2B**). MMP9 activity suggested by gel zymography was far stronger in venous compared with arterial thrombus. Utilizing cell surface biomarkers specific for leukocytes (CD45), thrombocytes (CD41) and erythrocytes (CD235a), immunoblotting revealed venous thrombi are more enriched than arterial thrombi in white blood cells (WBCs) and erythrocytes **(Figure 3A)**. The platelet content of venous and arterial thrombi was similar. The platelet and white blood cell content was similar in thrombi extracted from PA and the proximal venous circulation in the lower extremities. Interestingly, erythrocyte content was three-fold higher in thrombi extracted from the PA compared with that from the proximal venous circulation in the lower extremities (**Figure 3B-C**). Supporting in-gel zymography, immunoblotting thrombus extract for MMP isoforms revealed the thrombus contribution of MMPs overall was greater in venous compared with arterial thrombus and MMP9 was the dominant isoform expressed at the protein level (**Figure 3D**).

**Fig. 1B.**
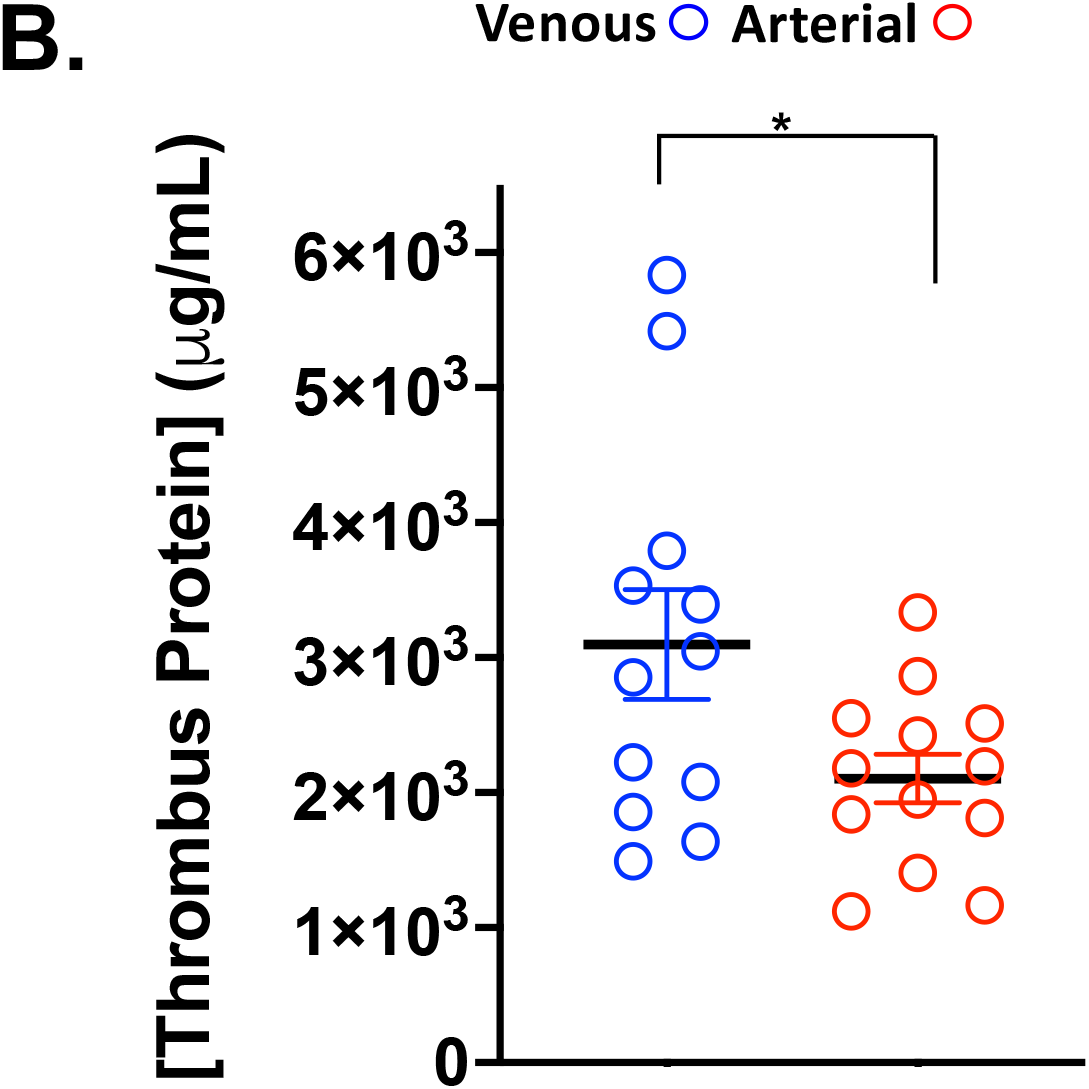
Thrombus from patients with thrombotic stroke and VTE: Arterial thrombi (n=13) and venous thrombi (DVT and PE, n=12) extracted by catheter and concentration of protein in 7 mg tissue was assessed. Venous thrombus was more heterogenous than arterial thrombus. Protein concentration was measured by the Bradford method and reported as mean ± SEM. *P=0.0311 between groups by Mann-Whitney *U*. VTE=venous thromboembolism.

**Fig. 2A.**
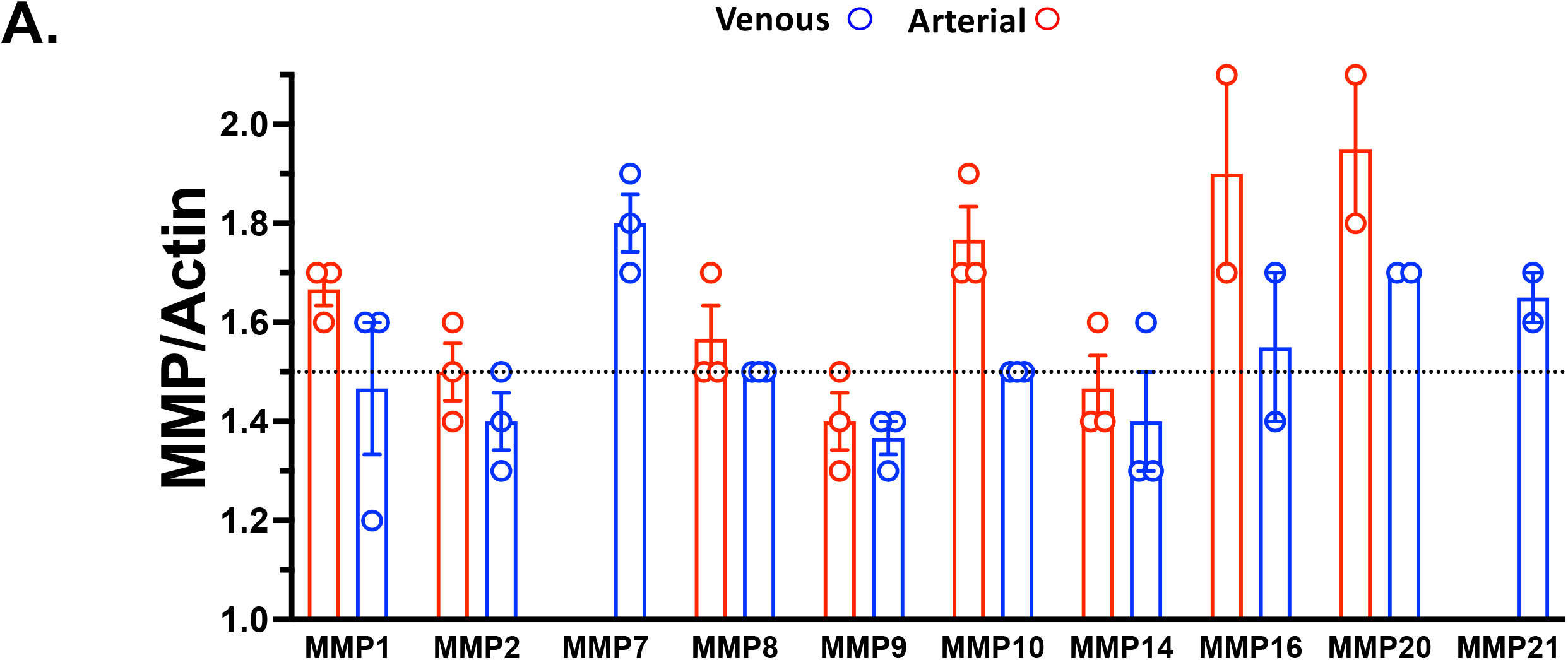
Matrix metalloproteinases presence in thrombus from patients with thrombotic stroke and VTE. RNA extracted from 7 mg thrombus was quantified by reverse transcriptase PCR (qRT-PCR) for Matrix metalloproteinases (MMPs). Where RNA was found in at least 3 thrombi from each group, the data were reported as mean ± SEM. The absence of data points indicates thrombus RNA (but not the positive control) was not detected.

**Fig. 2B.**
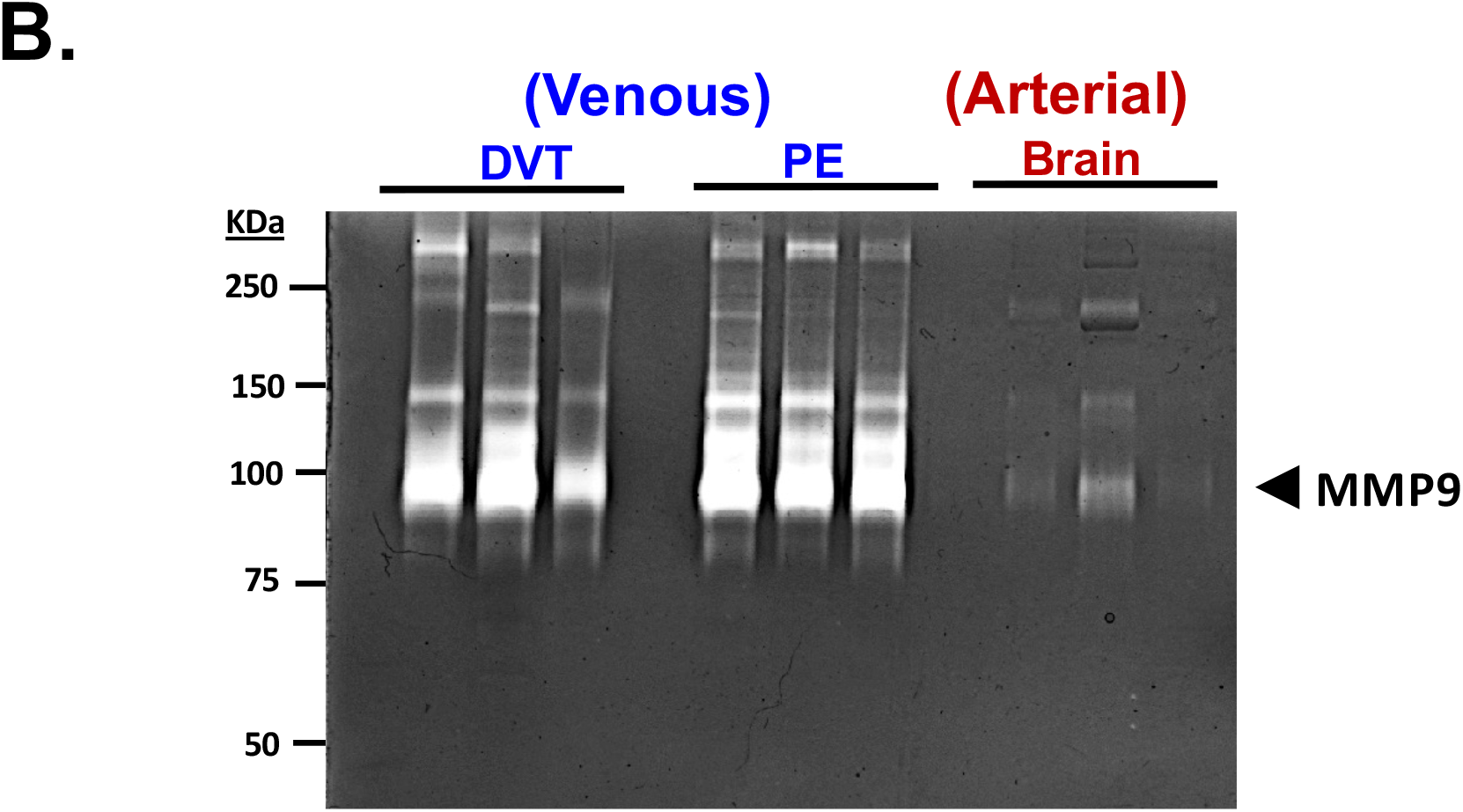
Matrix metalloproteinases activity in thrombus from patients with thrombotic stroke and VTE. Protein extracted from 7 mg thrombus was isolated and separated by non-reducing SDS-PAGE with gelatin embedded in the matrix. After washing, the gel was incubated in MMP reaction buffer overnight, and stained with Coomassie Blue reagent to highlight the presence of protein. A negative (white) impression indicates molecular weight regions where gelatin in digested by activated MMP gelatinase activity. MMP=matrix matolloproteinase. Molecular weight is indicated in KiloDaltons (KDa).

**Fig.3.**
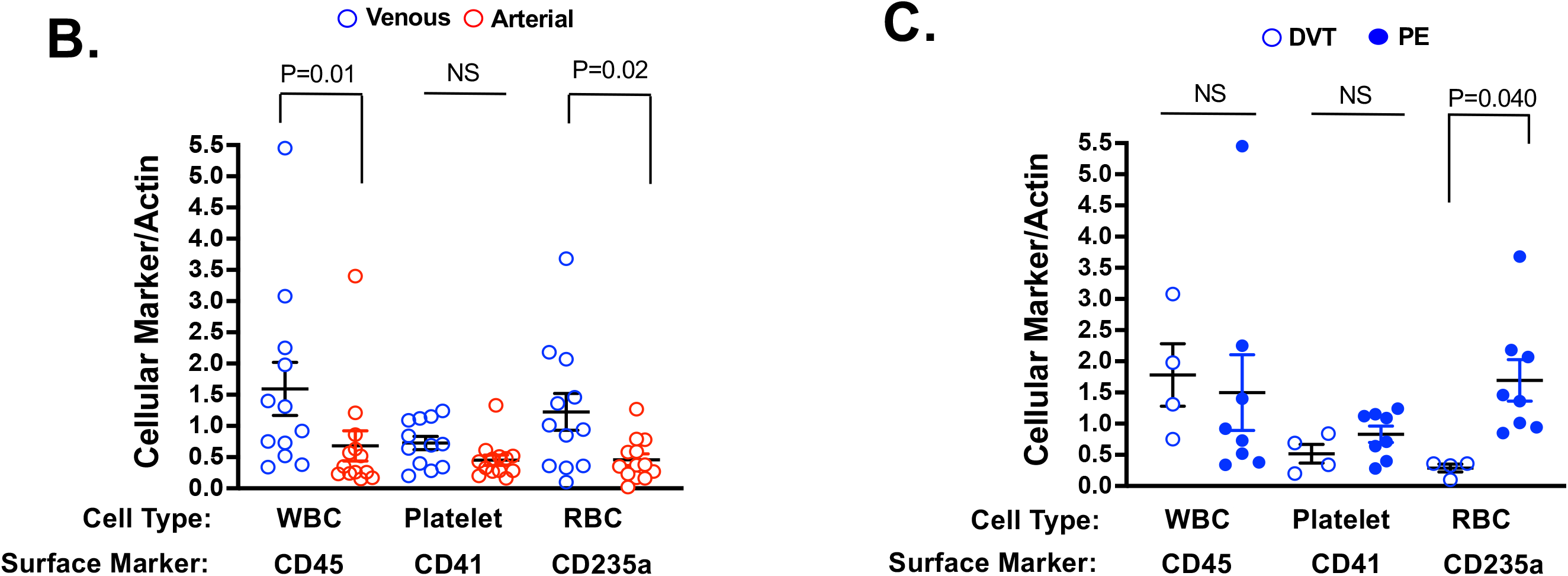
Circulating cellular markers in thrombus from patients with thrombotic stroke and VTE. **B.** Markers for white blood cells (WBC, CD45), platelets (CD41), and red blood cells (RBC, CD235a) in arterial and venous thrombi. **C.** Markers for white blood cells (WBC, CD45), platelets (CD41), and red blood cells (RBC, CD235a) in venous thrombi from the lower extremities and venous thromboembolic matter removed from the pulmonary artery. Protein extracted from 7 mg thrombus was isolated and separated by SDS-PAGE before immunoblotting using cellular protein markers as noted, and normalized to actin. Immunoreactive bands were quantified by densitometry and reported as mean ± SEM with a p value as noted. NS=not significant. VTE=venous thromboembolism. DVT=deep vein thrombosis. PE=pulmonary embolism.

**Fig. 3A.**
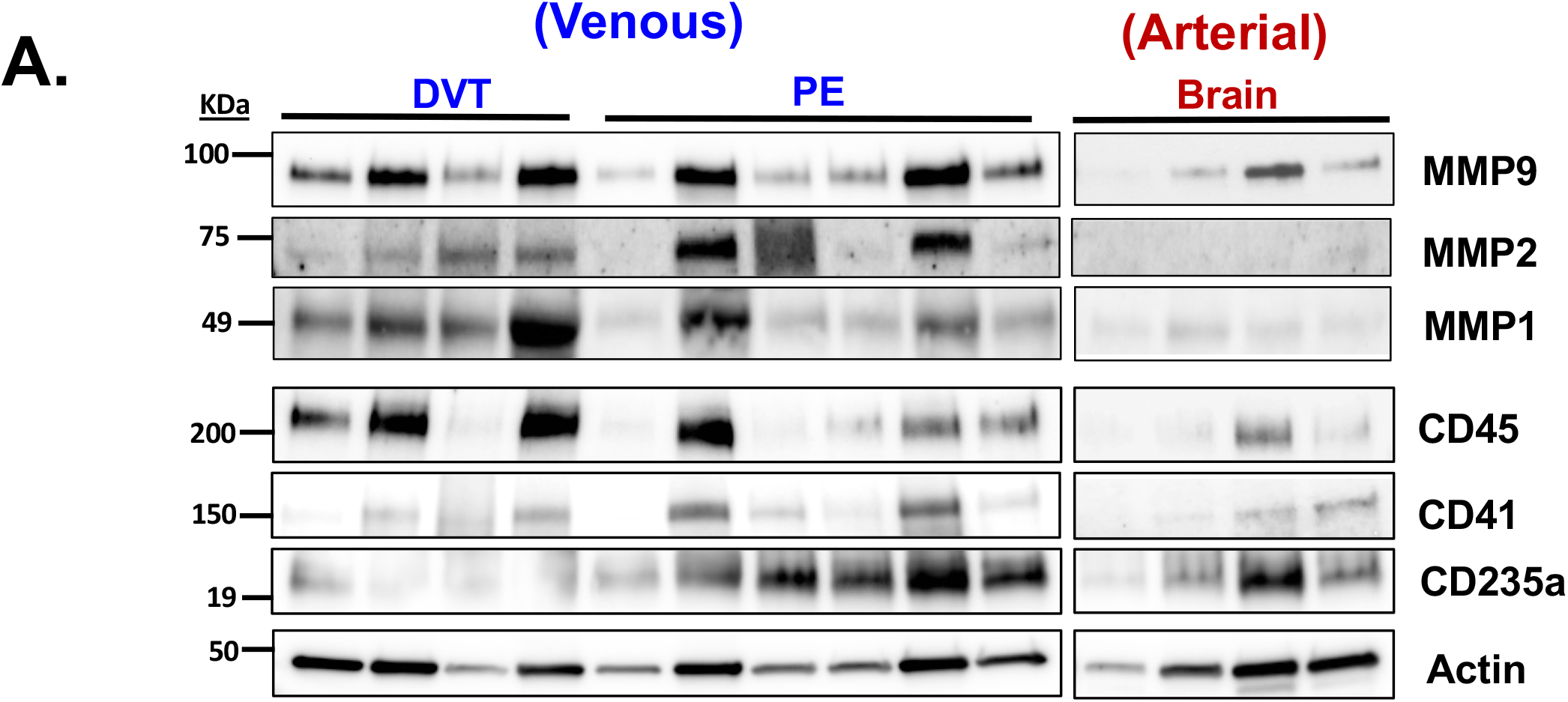
Circulating cellular markers in thrombus from patients with thrombotic stroke and VTE. Protein extracted from 7 mg thrombus was isolated and separated by SDS-PAGE before immunoblotting using protein markers found on white blood cells (WBC, CD45), platelets (CD41), and red blood cells (RBC, CD235a). Actin was used as a loading control. Immunoreactive bands were quantified by densitometry and reported as mean ± SEM. *P=0.01 (MWU), **P=0.02 (t-test), *** P=0.040 (MWU). MWU=Mann-Whitney *U*. NS=not significant. VTE=venous thromboembolism. DVT=deep vein thrombosis. PE=pulmonary embolism. Molecular weight is indicated in KiloDaltons (KDa).

**Fig. 3D.**
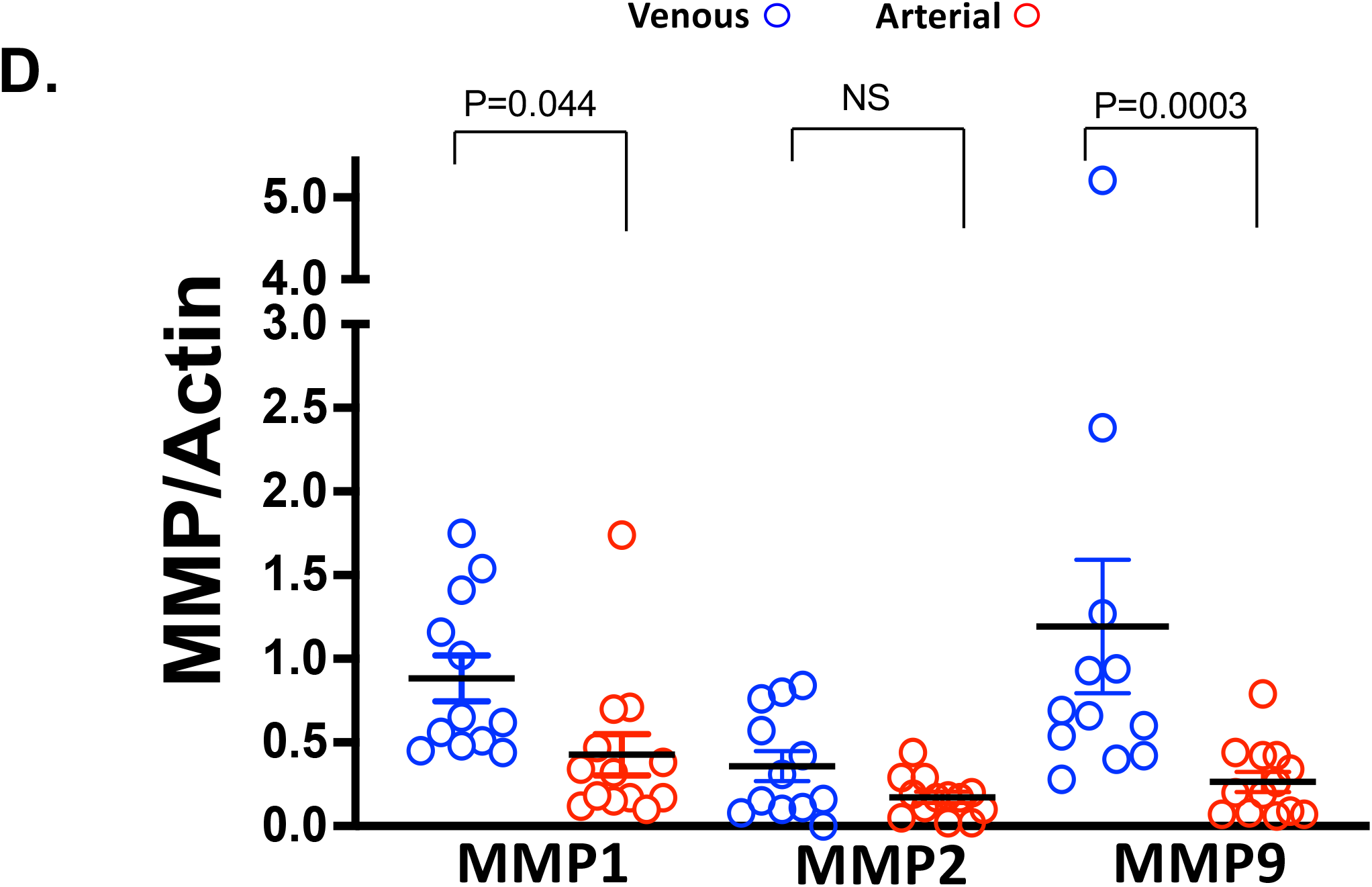
Matrix metalloproteinase content in thrombus from patients with thrombotic stroke and VTE. Protein extracted from 7 mg thrombus was isolated and analyzed for separated by SDS-Page then assessed for MMP1, MMP2, and MMP9 protein content. Actin is a loading control. Immunoreactive bands were quantified by densitometry and reported as mean ± SEM. P value as noted (MWU for MMP1 and MMP9 and t-testfor MMP2). MMP=matrix metalloproteinase.

### Molecular characteristics of thrombi from different vascular beds

RNA was extracted from various vascular beds to determine if the site of thrombus origin determines the thrombus phenotype. Capitalizing on the NanoString platform which is able to detect lower-yield or fragmented RNA common to aged tissue, sequence-specific mRNA probes were used to detect transcripts common to fibrotic and inflammatory pathways. Chronic thrombus abutting the vascular wall was collected from patients at the time of abdominal aortic aneurysm repair and used as a second source of arterial thrombus to compare to thrombus extracted from patients with acute CVA. Unexpectedly, the thrombus transcript expression profile in two distinct arterial beds (brain and aorta) was dissimilar, with only the anti-apoptotic marker *bcl-2* expression in aortic thrombus exceeding expression in brain thrombus (by five-fold). Pathways for organization of the extracellular matrix, fibrosis, adhesion, reflecting permanence in the arterial wall were upregulated and pathways for cellular communication and resolution were downregulated in older aortic thrombus compared with acute brain thrombus (**Figure 4A-C**). Similarly, comparing thrombus from the pre-capillary lung vasculature (pulmonary artery) to thrombus from the systemic arterial vasculature (brain), marked differences were noted in genes expression for angiogenesis, vascular inflammation and repair, apoptosis, and oxidative stress in the latter (**Figure 5A-C**). Comparing, acute lung thrombus (PA) to acute brain thrombus (CVA), marked differences in expression were noted with higher expression of endothelin and EGF signal transduction and B and T lymphocyte signaling transcripts in lung thrombus (**Supplemental Figures 1–2**). More intuitively, comparing thrombi from two different venous vascular beds (pulmonary artery and proximal veins of the lower extremity), fewer differences were noted in gene expression compared to thrombi between two different arterial beds. Gene expression for B lymphocyte and T lymphocyte signaling and inflammation were greater in DVT compared with PE thrombus, (**Supplemental Figures 3–4**).

**Fig. 4.**
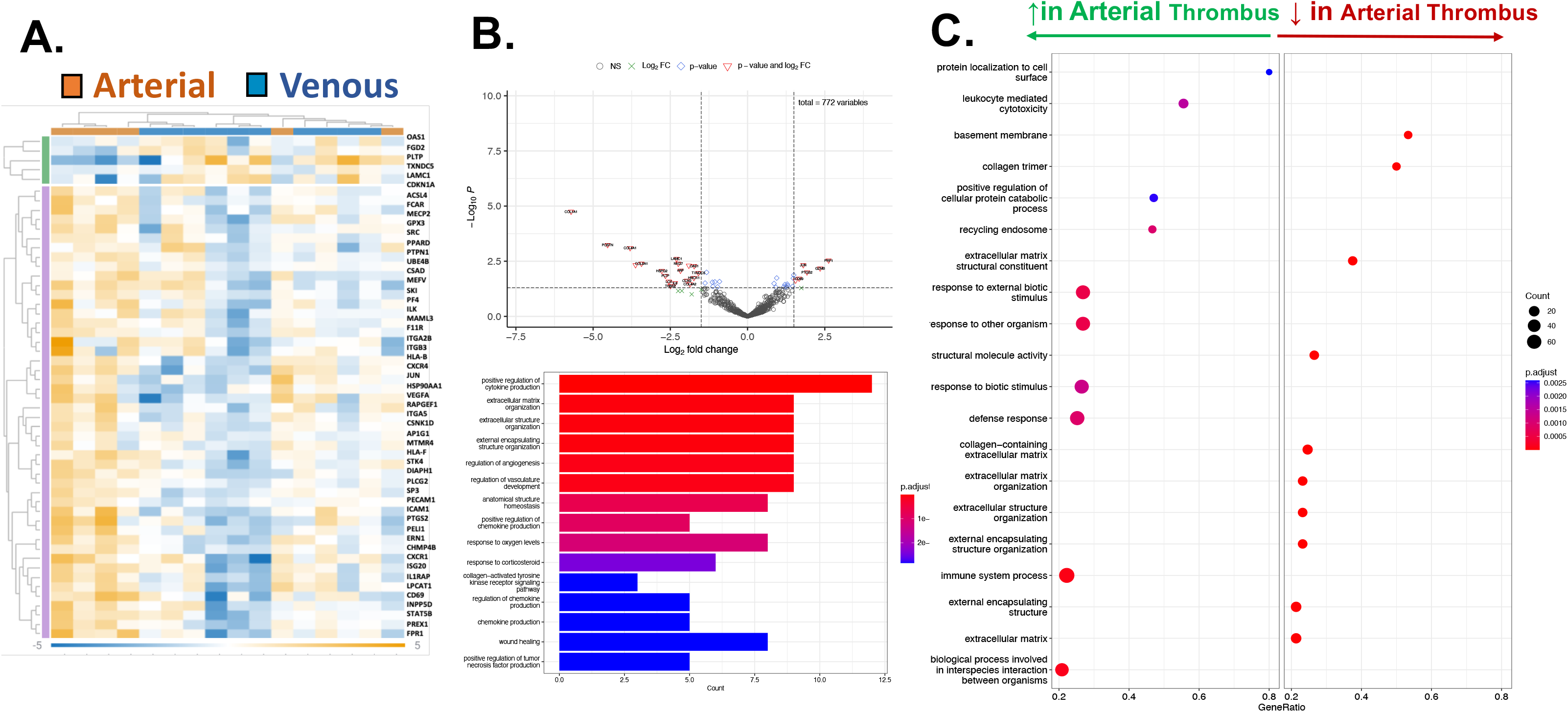
Transcriptomic analysis of arterial thrombus compared with venous thrombus. RNA was extracted and expression of genes involved in inflammation and fibrosis was determined in arterial thrombus extracted from patients with thrombotic stroke (n=6 cerebrovascular arterial) or venous thromboembolism (n=5 PE, n=5 DVT) at the time of catheter thrombectomy. **A.** Heat Map and **B.** Volcano plot and Gene Ontology analysis for biological pathways in the thrombus. **C.** Cellular and biochemical processes increased or decreased in arterial thrombus with respect to venous thrombus.

**Fig. 5.**
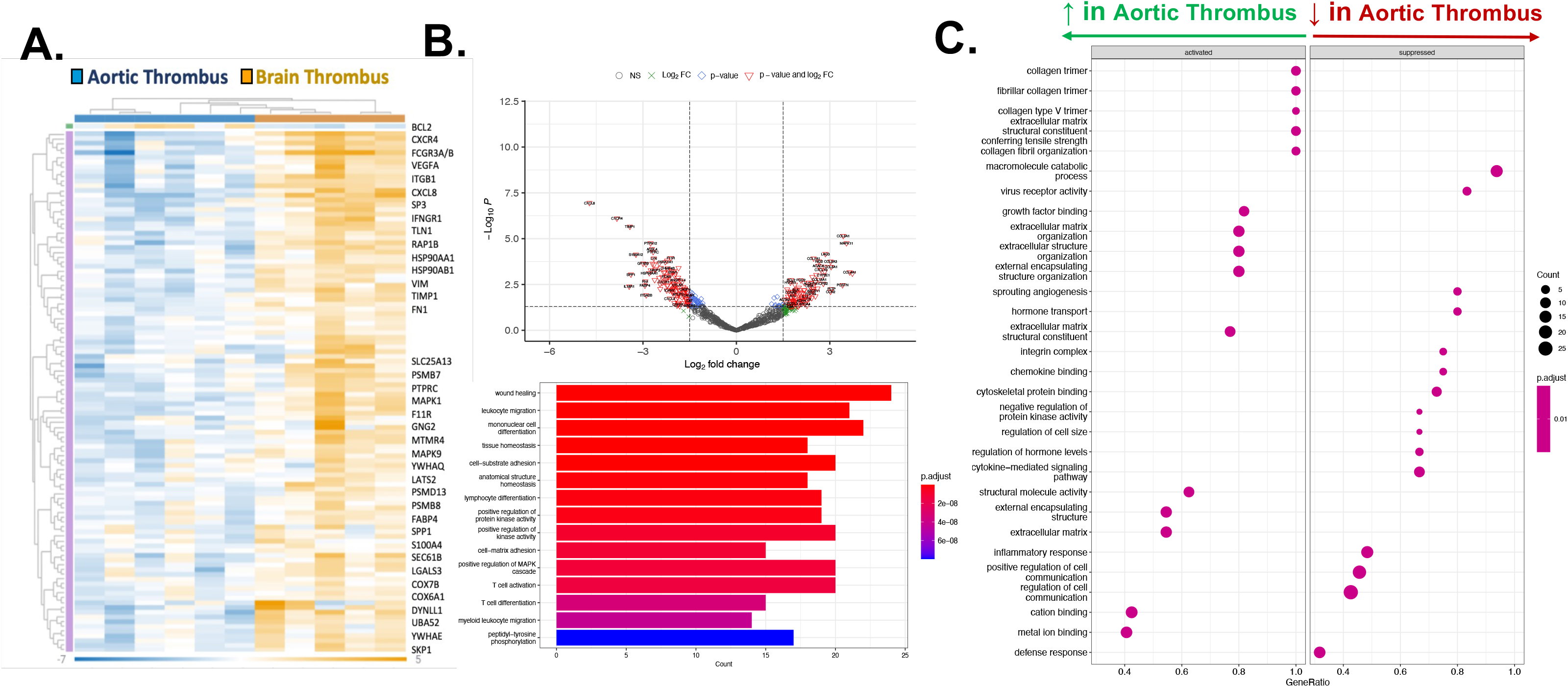
Transcriptomic analysis of arterial thrombus from different vascular beds. RNA was extracted and expression of genes involved in inflammation and fibrosis was determined in arterial thrombus extracted from patients with infrarenal AAA (n=6) at the time of surgical repair or from the brain following acute thrombotic stroke (n=5) at the time of catheter thrombectomy. **A.** Heat Map and **B.** Volcano plot and Gene Ontology analysis for biological pathways in the thrombus. **C.** Cellular and biochemical processes increased or decreased in aortic thrombus with respect to brain thrombus.

### MMP9 Enzymatic Activity in Arterial and Venous Thrombus

A MMP9-specific chromogenic substrate confirmed MMP9 specific activity is higher in venous than arterial thrombi (63 ± 8 ng/mL/μg protein vs. 25 ± 6 ng/mL/μg protein). (**Central Illustration**). MMP9 specific activity was significantly higher in catheter-extracted cerebrovascular thrombi in women compared with men (29.6 ± 6.3 ng/mL/μg protein vs. 10.4 ± 3.9 ng/mL/μg protein, p=0.020) and numerically greater but not reaching statistical significance in women compared to men with venous thrombi (63.96 ± 14 ng/mL/μg protein vs. 41.7 ± 14.0 ng/mL/μg protein, p=0.38) (**Supplemental Figure 5**).

## Discussion

The molecular and cellular composition of thrombus is disparate in four very different vascular beds: 1) the cerebral arterial circulation, 2) the aorta, 3) the pulmonary artery, and 4) the lower extremity proximal deep venous circulation. The most striking observation revealed in this investigation is that thrombus retains clear biological function, with MMP9 enzymatic activity directly proportional to the thrombus age and WBC content.

Consistent with prior studies, we observed increased WBC and RBC content in venous thrombi compared with arterial thrombi, especially those extracted from the pulmonary artery (26–28). Enriched WBC content represents a plausible source for the higher MMP9 activity observed in venous thrombus. While human platelets are known to secrete MMPs in specific cardiovascular pathologies (22,29,30), WBCs constitutively contribute most MMPs in the systemic circulation. Chernysh *et al*. used scanning electron microscopy to compare venous thrombus (DVT, PE) and arterial thrombus (coronary artery) and similarly found enriched RBC content in venous thrombus. Given similar platelet content in arterial and venous thrombi, the markedly higher MMP9 activity in venous thrombus is likely WBC-derived. In our data, venous thrombi were older than arterial thrombi by an order of 14-fold. Intuitively, the venous vasculature has lower sheer stress, allowing continuous cellular recruitment, and WBC enrichment as noted in venous thrombi.

While several MMP isoforms were detected in thrombus, MMP9, also known as gelatinase B, or Type IV collagenase, based on its ability to digest and penetrate the basement membrane (31), showed the greatest enrichment by enzymatic activity. Among the zinc-dependent metalloproteinases, MMP9 is known to remodel the extracellular matrix and tissues both in close proximity and distant to the origin of secretion. MMP9 has been implicated in infarct expansion and ventricular rupture after MI (32), in hemorrhagic transformation following CVA (33), and AAA growth, dissection, and rupture (23,34). This study is the first to identify active MMP9 in retrieved human thrombus specimens *ex vivo* following catheter extraction. Persistent thrombus abutting the blood vessel wall is a potential mediator of ongoing inflammation, scarring, and intimal angiogenesis preceding irreversible vascular damage and opposing thrombus resolution common to both PTS and CTED (35–37).

In relevant animal models of DVT, MMP9 is released into the circulation in early and intermediate periods of time post-acute venous thrombosis, then normalizes in the setting of thrombus resolution. Genetic MMP9 deletion prevents venous remodeling and biomechanical tissue dysfunction as well as permanent venous fibrosis (38–40). Furthermore, medications that suppress MMP9 activity *in vivo* in animals with experimental PE are protected from vascular remodeling and pulmonary hypertension (41). These mechanistic studies suggest incomplete thrombus resolution and continued MMP release may be factors that promote irreversible venous remodeling.

By their embolic nature and disparate origin of embolization, cerebral arterial thrombi intuitively have a more heterogeneous cellular composition than venous thrombi (42). Recent studies emphasize architectural patterns shared by thrombus in VTE and embolic CVA as well as the potential to evaluate thrombus-derived blood biomarkers to assist clinical decision-making including catheter thrombectomy, predicting thrombus susceptibility to thrombolytic drugs, and clinical prognostication (43,44). To further highlight the potential importance of thrombus-derived MMP not only as an adverse mediator of vascular remodeling, Zhong *et al*. elegantly show by way of a clinical study that blood MMP in the context of acute thrombotic CVA predicts short-term death and major disability (45). While we identified three-fold higher MMP activity in arterial thrombi extracted from women compared with men following acute CVA, the biological consequence of this remains unclear. Certainly, while the NIH stroke scale, time from symptoms to presentation, and use of intravascular thrombolysis are variables predicting hemorrhagic transformation, sex is not one of them (46). More broadly, however, the estrous cycle increases MMP9 activity in human cells (47), and MMP9 concentration appears to promote pathological vascular processes including arterial hypertension and vascular stiffening (48)

## Conclusions

While the CAVENT and ATTRACT trials disagree regarding the ability of catheter-directed thrombolysis to reduce the incidence of post-thrombotic syndrome in patients with DVT, a 15% decrease in the incidence of PTS after CDT at 24 months follow-up was observed (18,49). We provide mechanistic insight that thrombus composition and activity differ substantially based on the affected vascular bed and the duration of contact time between the thrombus and vessel wall. Focused translational research efforts on understanding how thrombus-derived mediators alter vascular function and thrombus resolution will be required to prevent often permanent symptoms of PTS and CTED.

### Limitations

This investigation, while conducted only in humans and using human tissues, is observational in nature, relying on patients’ recollection of symptom onset time. Furthermore, while blood vessels contain substrate for thrombus-derived enzymes, we do not directly demonstrate that MMP9 directly remodels blood vessels.

### Perspectives

#### Competency in Medical Knowledge

By performing genetic, biochemical, and cellular analyses of thrombus extracted from different vascular beds, this study revealed that thrombus age relates directly to its inflammatory and fibrotic capabilities.

#### Competency in Patient Care

##### Translational Outlook 1

Venous thrombi are not inert, but demonstrate considerable biological activity compared with arterial thrombi, secreting enzymes including MMP9 that can irreversibly remodel the blood vessel wall.

##### Translational Outlook 2

Cells enriched and continuously recruited to adherent thrombi in the vasculature are a source of secreted molecules that irreversibly remodel blood vessels, with a sex-dependence noted in arteries. Thrombus-derived blood biomarkers have probable utility in guiding decision-making for thrombectomy and should be prospectively evaluated.

## Abbreviations

DVT: Deep Vein Thrombosis
PE: Pulmonary Embolism
VTE: Venous Thromboembolism
CVA: Cerebrovascular Accident
CDT: Catheter-directed-therapy
MMP: Matrix Metalloproteinase

## Acknowledgements

The following grants were utilized for this study: HL158801 (to SJC). Nanostring assays were performed by the NIAID sponsored Clinical Trials in Organ Transplantation (CTOT) NanoString Core (U01 AI063594). We greatly thanks Karen Keslar for technical assistance and quality control. All authors have reported that they have no relationships relevant to the contents of this paper to disclose.

---

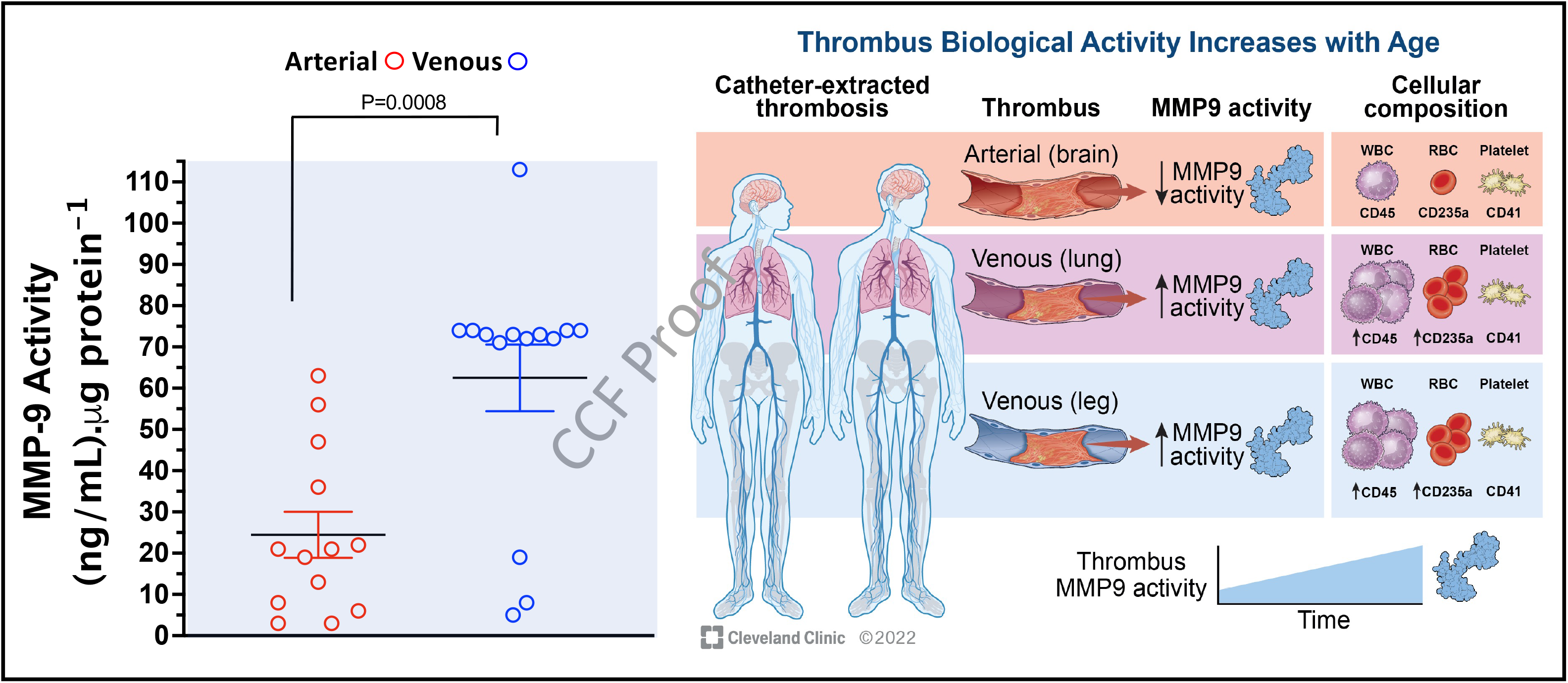
Central Illustration: Left: A MMP9-specific chromogenic substrate confirms MMP9 activity is markedly higher in venous than arterial thrombi (63 ± 8 ng/mL/μg protein vs. 25 ± 6 ng/mL/μg protein). Right: Venous thrombi are more enriched in inflammatory and fibrotic markers, WBC content, and MMP9 activity which are pathological signatures of chronic and irreversible remodeling of the blood vessel wall from older thrombus or later presentation.

## Supplemental Data

**Supplemental Figure 1.**
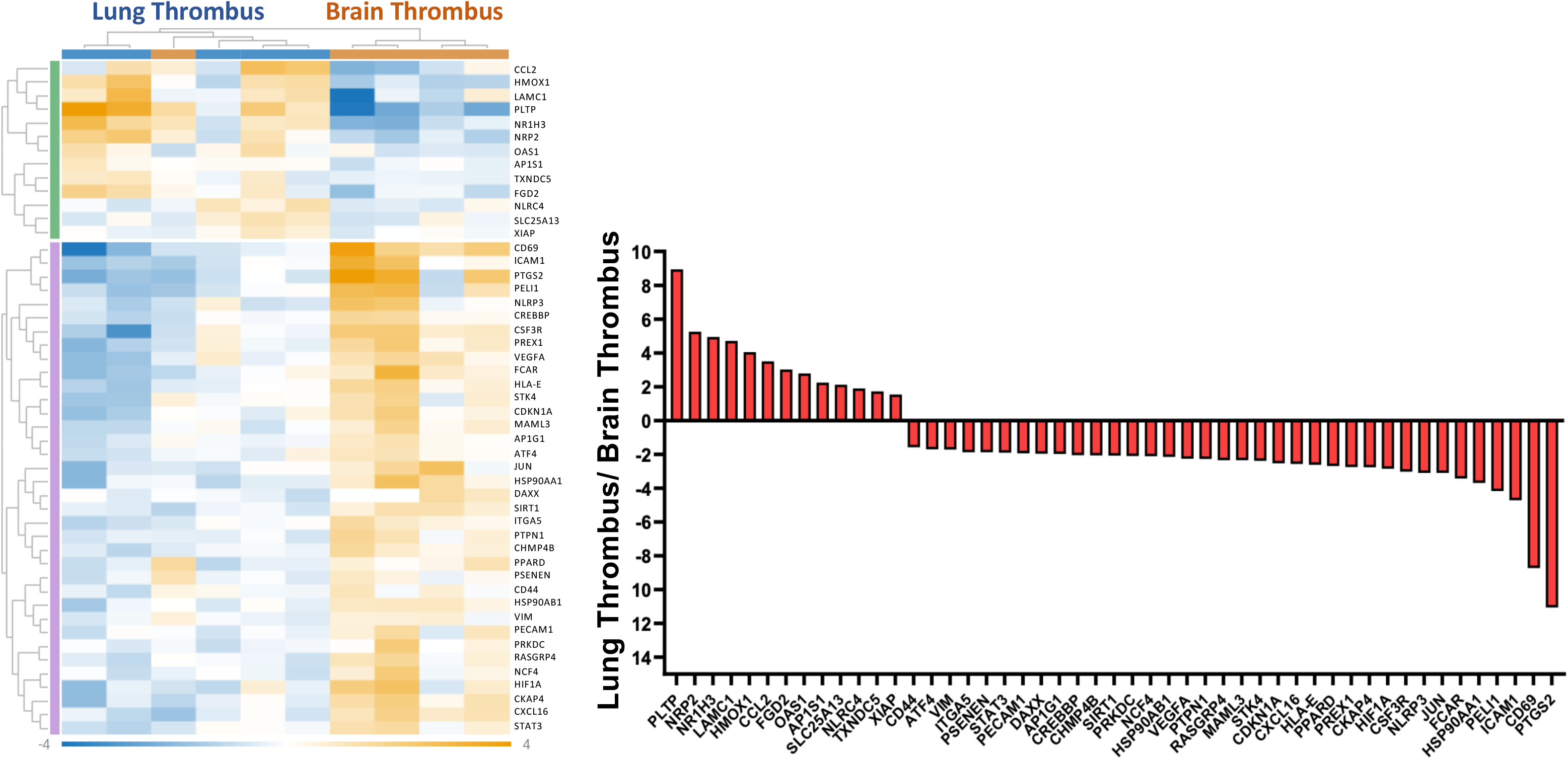
Transcriptomic analysis of lung thrombus compared to brain thrombus. RNA was extracted and expression of of genes involved in inflammation and fibrosis was determined in venous thrombus from patients with acute venous thromboembolism (n=5) following catheter thrombectomy or from the brain following acute thrombotic stroke (n=5) at the time of catheter thrombectomy.

**Supplemental Figure 2.**
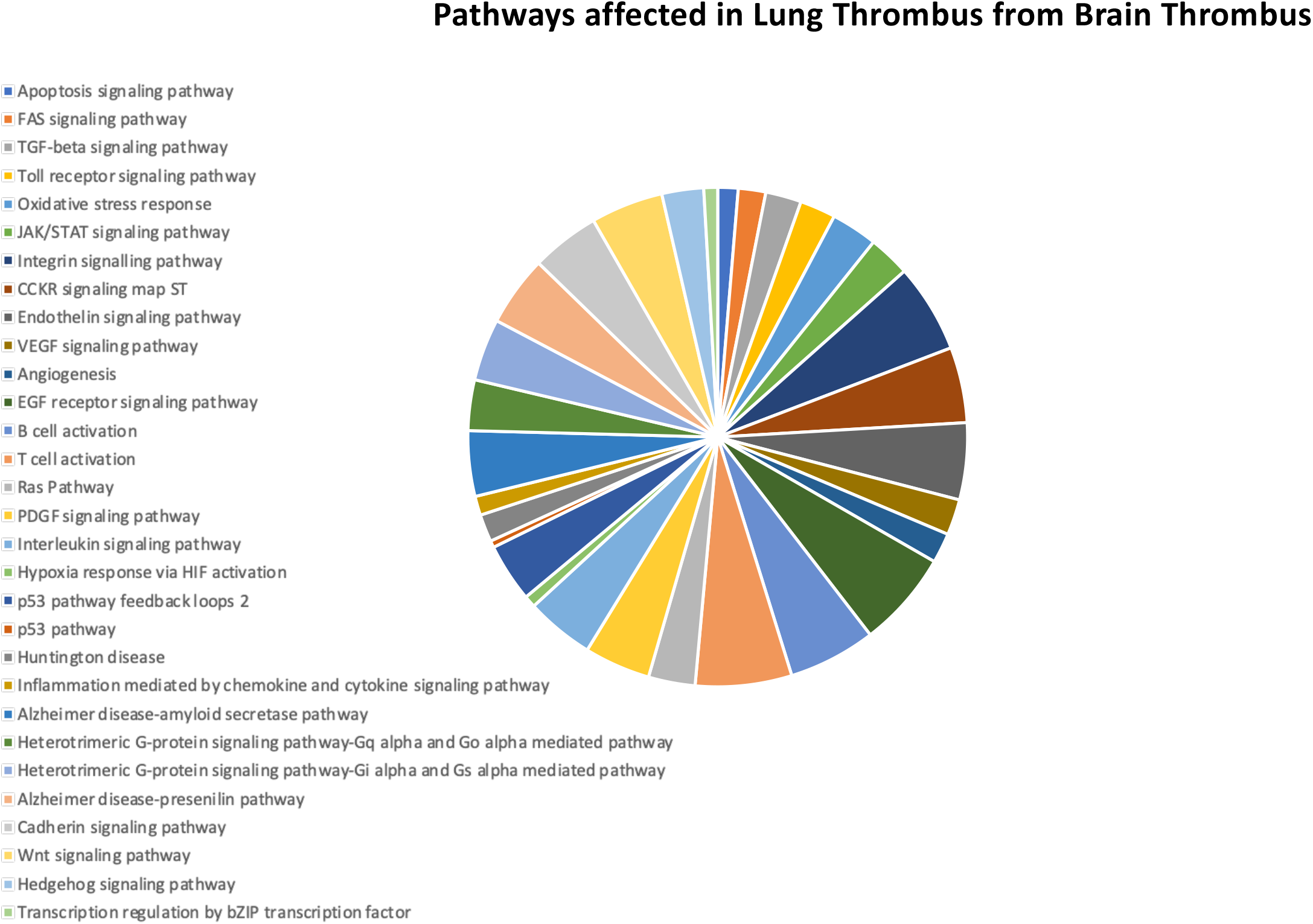
Pathways analysis by NanoString for biological pathways that differ in lung thrombus (numerator) compared with brain thrombus (denominator).

**Supplemental Figure 3.**
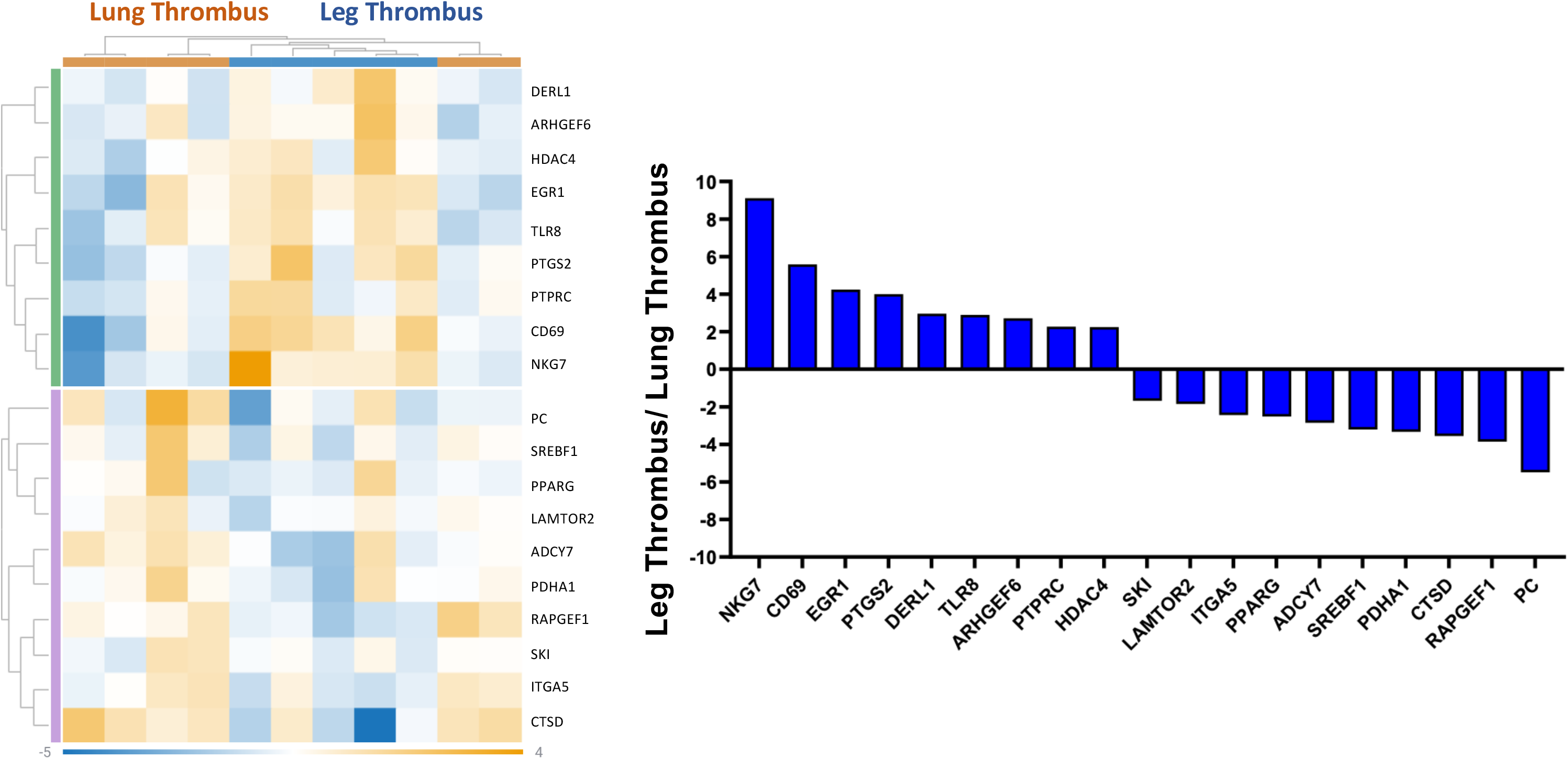
Heat map (left) and comparative expression of major gene (right) by NanoString that differ in leg thrombus (numerator, n=6) compared with lung thrombus (denominator, n=5). Filter is −1.5 to 1.5 p=0.05

**Supplemental Figure 4.**
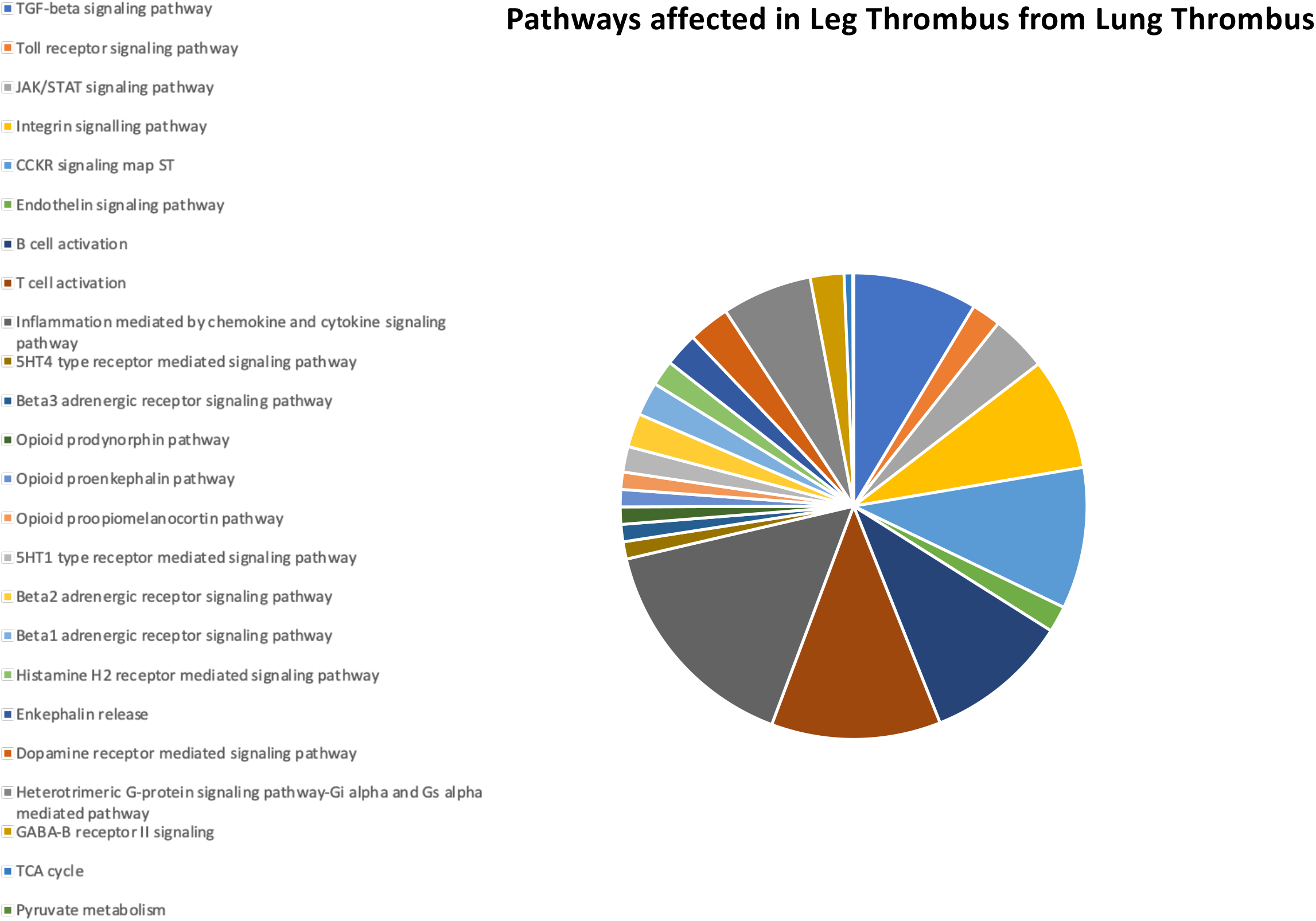
Pathways analysis for biological pathways by NanoString that differ in leg thrombus (numerator) compared with lung thrombus (denominator).

**Supplemental Figure 5:**
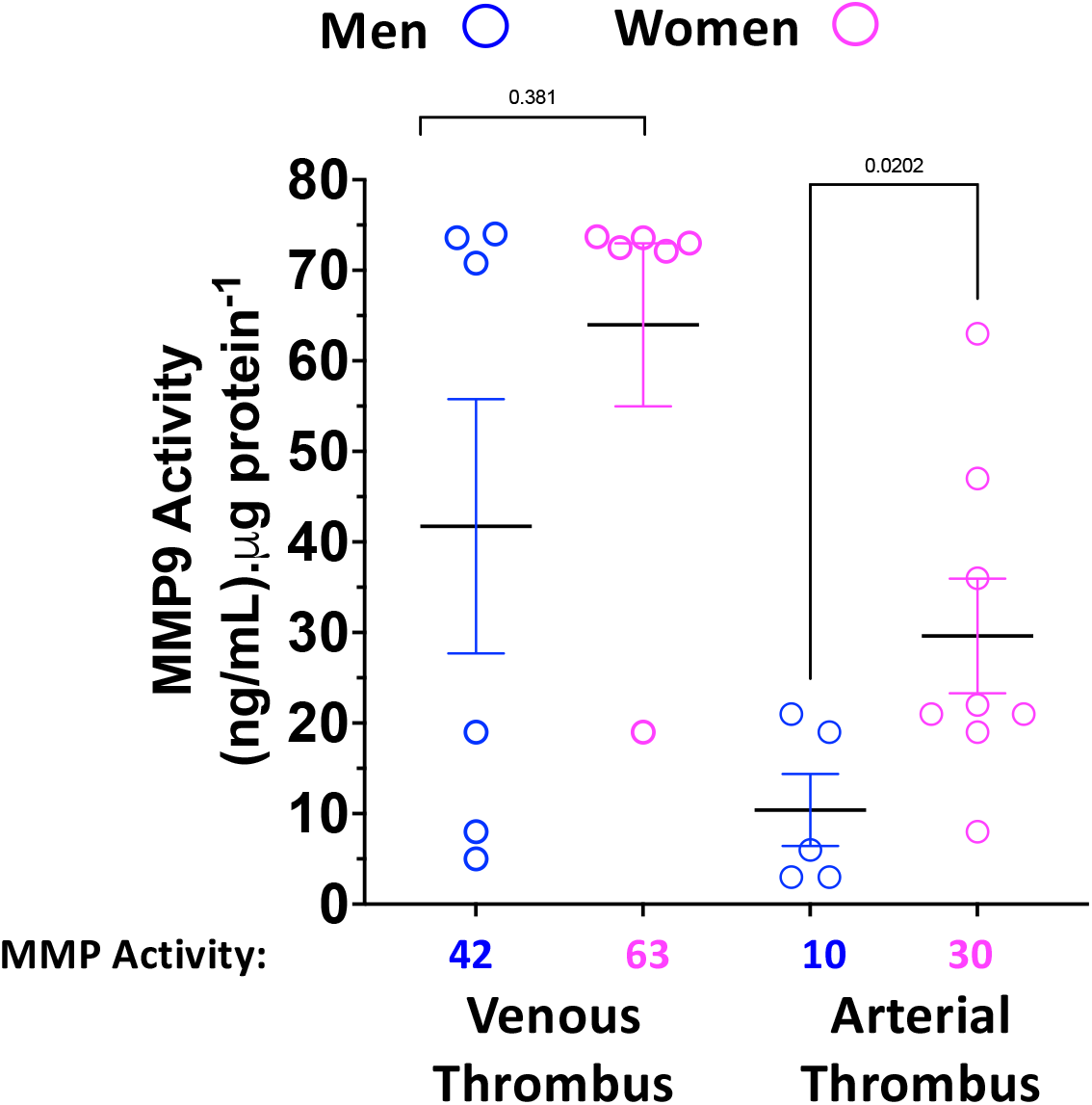
Thrombus MMP9 activity as a function of sex for arterial thrombus (n=13; 5 men/8 women) and venous thrombus (n=12; 6 men / 6 women) using a chromogenic substrate and represented as mean activity ± SEM (numerical data below graph). P value as noted between groups (MWU). MMP=matrix metalloproteinase.

